# Incorporation of unnatural amino acid for the tagging of cannabinoid receptors 1 and 2 reveals receptor roles in regulating cAMP levels

**DOI:** 10.1101/2021.06.09.447758

**Authors:** Alix Thomas, Carsten Schultz, Aurélien Laguerre

**Affiliations:** Dept. of Chemical Physiology & Biochemistry, OHSU, Portland, OR, USA

## Abstract

The role of CB1/CB2 co-expression in cell signaling remains elusive. We established a simplified mammalian cell model system in which expression of CB1 or CB2 can be easily monitored under a confocal microscope. For this, we applied amber codon suppression in live cells to incorporate a single *trans*-cyclooctene (TCO) bearing amino acid in one of the extracellular loops of CB1 or CB2, followed by fluorescent labeling *via* click chemistry. We employed genetically encoded biosensors to measure the roles of CB1 and/or CB2 in regulating intracellular calcium ([Ca^2+^]_i_) and cAMP ([cAMP]_i_) levels. We show that the agonist-mediated activation of tagged-CB1 or -CB2 can transiently elevate [Ca^2+^]_i_ levels. However, when the two receptors were co-expressed in the same cell, CB2 no longer signaled through calcium although CB1-mediated transient elevation of [Ca^2+^]_i_ levels was unaffected. Because of the existence of crosstalk between calcium and cAMP signaling, we measured the effects of CB1 and/or CB2 in regulating adenylate cyclase activity. We found that the expression of CB1 increased forskolin-induced [cAMP]_i_ levels compared to non-transfected cells. Conversely, CB2 expression decreased stimulated [cAMP]_i_ levels under the same conditions. Finally, co-expressed CB1 and CB2 receptors showed additive yet opposing effects on stimulated [cAMP]_i_ levels. These observations suggest that co-expressed CB1/CB2 act locally as a pair in regulating cell excitability by modulating stimulated [cAMP]_i_ levels.

## Introduction

Cannabinoid receptors 1 and 2 (CB1 and CB2) belong to the subfamily of class-A rhodopsin-like GPCRs.^1,2^ In mammals, CB1 and CB2 frequently co-express in central and peripheral tissues,^3,4^ where the two receptors are co-activated by the endocannabinoid 2-arachidonoylglycerol (2-AG) –full agonist of both receptors.^5^ However, the functional consequences of CB1/CB2 co-expression have not yet been investigated in detail. CB1 and CB2 are individually known to play fundamental roles in membrane plasticity,^6^ vesicle secretion,^7,8^ cell migration^9^ and inflammation.^10^ CB1 and CB2 share only 44 % structural homology^11,12^ but both receptors predominantly signal via Gi/o proteins that ultimately lead to changes in intracellular calcium and [cAMP]_i_ levels.^13,14,15^ CB-mediated fluctuations in [cAMP]_i_ levels connect to the regulation of a variety of targets such as A type and inwardly rectifying potassium channels and focal adhesion kinase.^16,17^ CB1 and CB2 also signal via Gβγ subunit that transduces receptor activation to phosphatidylinositol 3-kinase (PI3K) and mitogen-activated protein kinases.^18^

Despite using similar signaling components, the pharmacological interference with CB1 or CB2 has suggested opposing roles of the two receptors. In the central system, opposing roles for CB1 and CB2 receptors were recently reported in modulating behavioral responses to cocaine.^19^ Here, the CB1 reverse agonist rimonabant prevented the increase in c-Fos expression induced by cocaine on the shell region of the nucleus accubens in the brain, an effect that can be reversed by treatment with the CB2 reverse agonist AM630. In the immune system, the presence of both CB1 and CB2 suggests that the two GPCRs play a role as immunomodulators.^20^ While CB1 prolonged activation is often associated with an increase in inflammatory response,^21^ CB2 has significant anti-inflammatory functions.^22,23,24^ Co-expression of mRNA for both CB1 and CB2 has been documented in the murine spleen as well as in brain-resident macrophage-like microglia.^25^ Furthermore, mRNA and protein expression corresponding to both CB1 and CB2 have recently been found in two mast cell lines.^26^ In this study, the authors showed that co-expressed receptors were not functionally redundant, but were having distinct functions. In the cardiovascular system, CB1 and CB2 were also found to act in opposing ways under pathological conditions. In patients suffering from heart failure, elevated peripheral blood levels of endocannabinoids were measured.^27^ Moreover, whereas the expression of CB1 receptors was downregulated 0.7-fold, CB2 receptors were upregulated more than 11-fold, indicating a shift toward CB2 expression in cardiomyocytes of the human myocardium.^28^ Another example can be found in the pancreas where insulin is secreted in a pulsatile fashion by β-cells.^29^ The amount of hormone released in the extracellular space often correlates with an increased frequency of [Ca^2+^]_i_ oscillations in the secreting cells.^30^ We and others have shown that CB1 and CB2 receptors have counteracting effects on insulin secretion.^31,32,33^ By using a photo-activatable (caged) 2-AG,^34^ we demonstrated that CB1 and CB2 were activated by the endocannabinoid in a concerted manner that regulated intracellular [Ca^2+^]_i_ levels.^31^ We further showed that while CB1 activation was required to maintain intracellular [Ca^2+^]_i_ oscillations, CB2 activation had a dampening role.

The combined observations suggest that co-expressed CB1 and CB2 receptors are juxtaposed in a way that one’s activation might regulate the other’s activity in many cell types. Compounds aiming at CB1 or CB2 have been used or studied as treatment for pain, seizures, psychiatric disorders, obesity, diabetes, metabolic diseases, neurodegenerative diseases, and cancer.^35^ Maybe not surprisingly, these attempts have not resulted in novel treatment –as the system is not modulated as a whole. Investigating the functions of CB1 and CB2 as a pair rather than as single targets could spruce the development of new therapeutic strategies. It is therefore of great interest to determine the biological function of CB1/CB2 co-expression and their interplay in regulating cell signaling. An intuitive step towards this objective is the development of a simplified system to study cannabinoid signaling.

HeLa cells are devoid of endogenous CB1 or CB2 and transcriptome and immunohistochemistry analysis showed the absence of CB1 and CB2 but demonstrated that Gi and Go are abundant (Figure S1A). CB1 and CB2 are both fully activated by the endocannabinoid 2-AG which is biosynthesized via the action of diacylglycerol lipase α (DAGLα) and is further degraded by the monoacylglycerol lipase MAGL.^36^ Interestingly, HeLa cells express both DAGLα and MAGL (Figure S1A). Co-expressing CB1 or CB2 in HeLa cells therefore formed the starting point for a minimal cannabinoid signaling system in a mammalian cell line. Fluorescent fusion proteins (FPs) are commonly used to investigate the location and trafficking of GPCRs. However, examples have shown that FPs can sometimes interfere with the receptor’s activity and complicate the interpretation of experimental observations.^37^ To avoid such interference, alternative tagging approaches using single non-canonical amino acids and click chemistry are an attractive option. In this study, amber stop-codon suppression^38^ was successfully applied in live mammalian cells to incorporate a single *trans*-cyclooctene (TCO) bearing amino acid in one of the extracellular loops of CB1 and CB2, respectively. The tagged receptors were successfully used to assess the roles of CB1 or CB2 in regulating [Ca^2+^]_i_ and cAMP [cAMP]_i_ levels.

## Results

Immunofluorescence experiments demonstrated the endogenous expression of CB1 and CB2 in the positive control MIN6 β-cells but did not detect the presence of endogenous cannabinoid receptors in HeLa Kyoto cells (Figure 1A). We therefore chose these cells as a model for reconstituting the CB1/CB2 switch in a benign host cell. Without expressing CB1 or CB2 in HeLa cells, we did not observe an effect of the unspecific agonist WIN55,212-2 on intracellular calcium levels measured by the genetically encoded indicator R-GECO, although ATP addition (100 μM) induced a major calcium transient, (F/F_0_=3.339±0.119, n=89) due to its binding and signaling via the Gq/11 coupled P2Y receptor (Figure 1B).^39^ P2Y and Gi, Go, and Gq have high transcriptional levels in Hela cells suggesting a high level of expression. (Figure S1). Accordingly, inhibition of phospholipase C (PLC) downstream of Gq/11 by U73122 (10 μM) significantly reduced the cell response to ATP (F/F_0_=1.528±0.066, n=99) while the protein kinase A (PKA) inhibitor H89 had little effect (F/F_0_=2.838±0.116, n=114, Figure 1E). Note that U73122 is also known to inhibit the calcium transient evoked upon photolysis of caged IP_3_ as well as the direct activation of the ryanodine receptor with caffeine, which both do not require PLC activation.^40^

**Figure 1.**
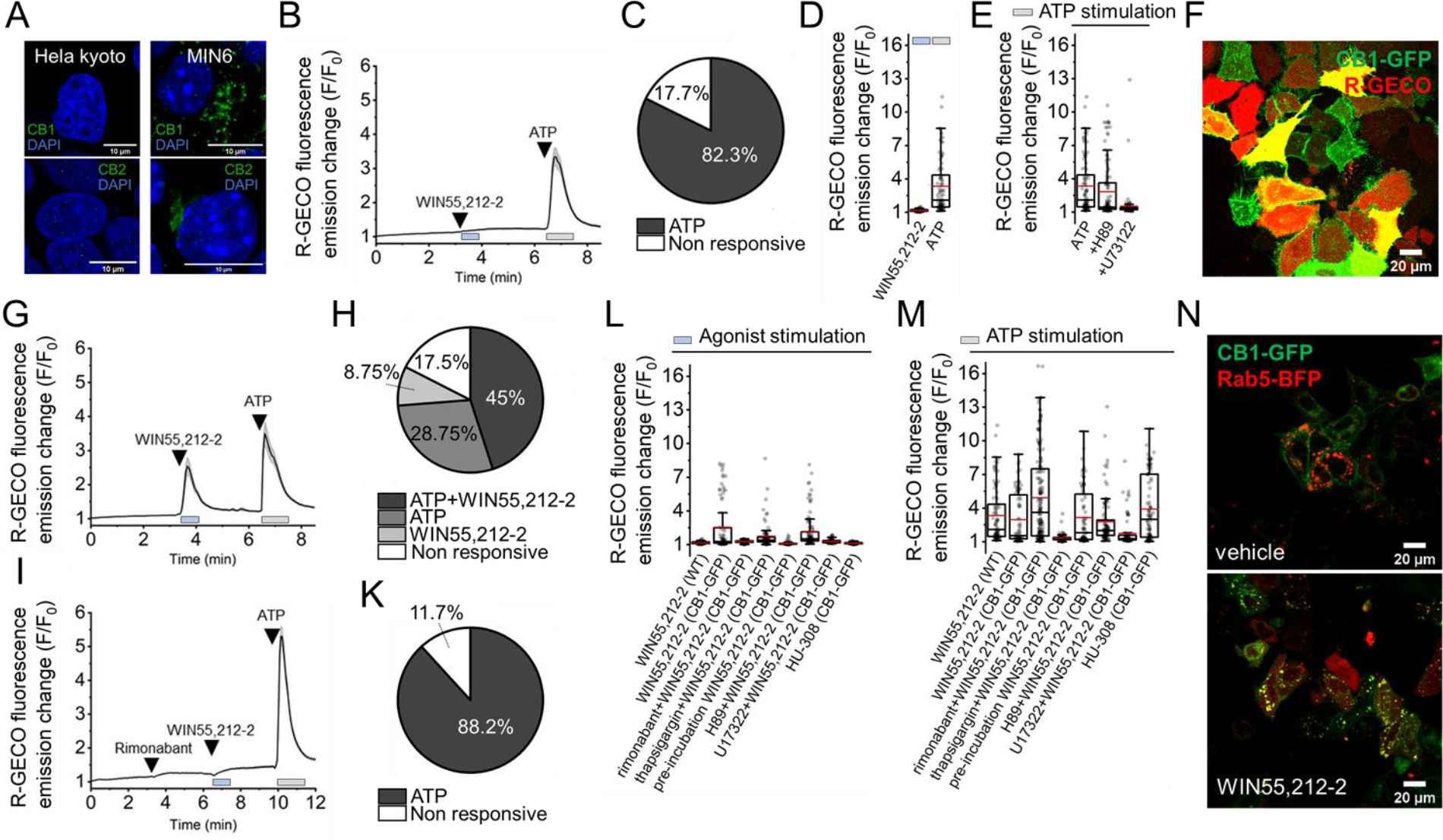
**A.** Immuno-fluorescence experiments comparing CB1 and CB2 expressions in HeLa Kyoto and in MIN6. **B.** Fluctuations of [Ca^2+^]_i_ in wild type HeLa Kyoto subjected to treatment with the CB1/CB2 agonist WIN55,212-2 (10 μM) and with ATP (100 μM). **C.** Corresponding pie chart showing the distribution of ATP-responsive versus non-responsive cells. **D.** Quantification of maximum R-GECO fluorescence emission intensity F/F0 after stimulating the cells with WIN55,212-2 (blue square) or ATP (grey square). Red line represents mean value. **E.** ATP-mediated calcium transient in wild type HeLa Kyoto alone, or in presence of H89 or U73122. **F.** Confocal micrograph showing the co-expression of CB1-GFP and the calcium sensor R-GECO. **G.** Fluctuations of [Ca^2+^]_i_ in CB1-GFP-transfected HeLa Kyoto subjected to treatment with the CB1/CB2 agonist WIN55,212-2 (10 μM) and with ATP (100 μM). **H.** Pie-chart showing the repartition of the different cell populations responding to treatments with CB1 agonist and ATP on CB1-GFP transfected HeLa Kyoto cells. **I.** Same experiment performed in the presence of CB1 inverse agonist rimonabant (10 μM). **K.** Pie-chart showing the repartition of the different cell populations responding to treatments with CB1 agonist and ATP in the presence of rimonabant on CB1-GFP transfected HeLa Kyoto cells. **L.** Comparison of maximum fluorescence increase of R-GECO in CB1-GFP-transfected HeLa Kyoto after treatment with WIN55,212-2 (10 μM) under different experimental conditions. **M.** Comparison of maximum fluorescence increase after treatment with ATP (100 μM) under different experimental conditions. **N.** Confocal micrographs showing the relative subcellular locations of CB1-GFP and Rab5-BFP with vehicle or with WIN55,212-2 (10 μM) for 3 hr.

We then investigated the effects of CB1 expression and activation on calcium levels. We initially expressed a version of CB1 fused to GFP as well as the calcium indicator R-GECO (Figure 1F).^41^ We employed a semi-automatized cell segmentation method allowing us to extract single calcium traces from all the cells present in the microscope’s field of view (Figure S1B). This approach permitted to sort and quantify different cell populations depending on their response profile (*i.e.*, either the cell responds to WIN55,212-2 or ATP only, responds to both compounds or none of them, Figure S1C to I). As expected, expression of CB1-GFP and subsequent activation with WIN55,212-2 quickly triggered a transient increase in [Ca^2+^]_i_ levels (F/F_0_=2.545±0.128, n=80, Figure 1G). This CB1-mediated calcium transient was completely abolished by pre-treating the cells with the specific CB1 inverse agonist rimonabant (F/F_0_=1.267±0.003, n=170, Figure 1I). Intracellular calcium stores are known to be involved in the CB1-mediated increase in cytoplasmic calcium levels in different cellular models.^42,43,44,45,46^ This was confirmed by a short incubation with thapsigargin, a non-competitive inhibitor of the sarco/endoplasmic reticulum Ca^2+^ ATPase (SERCA), that reduced the calcium response to WIN55,212-2 (F/F_0_=1.761±0.131, n=80, Figure 1I). Chronic activation of CB1 is known to lead to receptor internalization and subsequent desensitization. Therefore, we co-expressed CB1-GFP together with the early endosome marker Rab5-BFP and monitored protein co-localization in presence or absence of the CB1 agonist (Figure 1N, S1K and S1L). In the presence of WIN55,212 for 3 hr, we found prolonged activation of CB1 triggered receptor internalization and co-localization with Rab5 in early endosomes. In addition, a second dose of the agonist was unable to provide another transient (F/F_0_=1.107±0.008, n=80, Figure 1M) indicating that the receptor was desensitized upon internalization. As an additional control, we showed that the prolonged inactivation of CB1 with the reverse agonist rimonabant (10 μM) for 3 hr lead to the accumulation of the receptor at the plasma membrane (Figure S1L).

A recent study demonstrated that CB1-mediated calcium increase was reduced by blocking nicotinic acid-adenine dinucleotide phosphate- or inositol 1,4,5-trisphosphate-dependent calcium release, which are known to release calcium from the lysosome and the endoplasmic reticulum (ER), respectively.^47^ Furthermore, the authors showed that the transient was completely abolished when both lysosomal and ER calcium release pathways were blocked. In our HeLa cell system, U73122 and thapsigargin significantly reduced the CB1-mediated calcium transient confirming the involvement of IP_3_ receptors in CB1-mediated calcium release (Figure 1L, F/F_0_=1.288±0.009, n=87 and F/F_0_=1.761±0.065, n=80, respectively). As an additional control, we showed that PKA inhibition with H89 did not impede the CB1 signaling on calcium (F/F_0_=2.153±0.077, n=97). Finally, we showed that the specific CB2 agonist HU308 was ineffective in CB1 transfected cells (Figure 1L, F/F_0_=1.132±0.004, n=96). While the presence of SERCA or PLC inhibitors blocked the ATP-mediated calcium transient (F/F_0_=1.335±0.387, n=80 and F/F_0_=1.761±0.136, n=87, respectively), the expression and subsequent activation of CB1 did not affect the cell response to ATP (F/F_0_=2.981±0.125, n=80). However, quite unexpectedly, we observed that the CB1-specific inverse agonist rimonabant significantly enhanced ATP-stimulated calcium transients (F/F_0_=4.936±0.151, n=170, Figure 1M).

To ensure that the steric perturbations engendered by the fused GFP did not interfere with cannabinoid receptor signaling on calcium, we reduced the size of the tag to a single amino acid and compared the effects of both modified receptors on the activation of [Ca^2+^]_i_ levels (Figure 2). We incorporated *trans*-cyclooct-2-en-L-lysine (TCO*A) for permitting catalyst-free ultrafast labeling of the receptors, as previously described for other membrane proteins.^48,49^ We employed the orthogonal tRNA/tRNA synthetase pair (tRNA/RS) from *Methanosarcina mazei* to introduce TCO*A to the first extracellular loop of CB1 and on the N-terminal tail of CB2, respectively. Of several positions examined, TCO*A incorporation and expression levels were most successful by replacing phenylalanine at position 180 in CB1 and Ser29 in CB2 (see Table S1 for details). CB1-F180 or CB2-S29 allowed post-translational tagging of the proteins with a methyl tetrazine-bearing dye for visualizing the receptors in living cells, likely with minimal structural impact. For labeling before calcium imaging experiments, we choose the cell permeable dye methyl tetrazine ATTO 488 (ATTO 488 MeTet, Figure 2A and B). Activating CB1-F180 with WIN55,212-2 transiently increased [Ca^2+^]_i_ levels in the same order of magnitude as in the case of CB1-GFP expression. The response to WIN55,212-2 was also blocked by the CB1 inverse agonist rimonabant (from F/F_0_=2.531±0.111, n=147 versus F/F_0_=1.267±0.01, n=145, Figure 2C). Similar to what we described above, the treatment with rimonabant had a positive effect on the ATP-mediated calcium transient (F/F_0_=4.336±0.162, n=145) and thapsigargin drastically reduced the effects of CB1-F180 activation on intracellular calcium levels (F/F_0_=1.510±0.312, n=51). Calcium-free media conditions led to a significant reduction in the size of the CB1-mediated calcium transient but did not block the response fully (F/F_0_=1.762±0.129), indicating again that release from intracellular stores drives the calcium signal.

**Figure 2.**
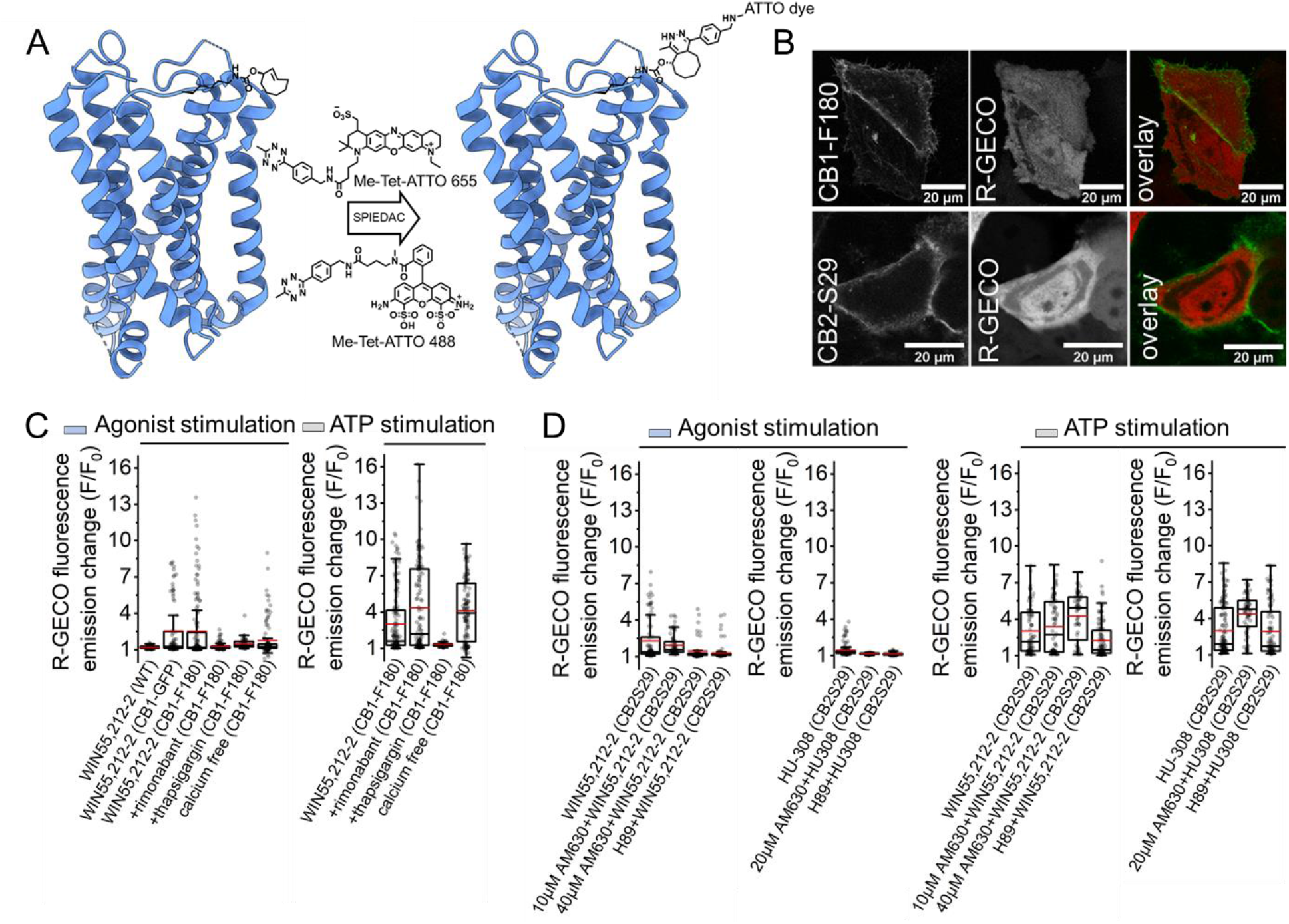
**A.** Illustration of cannabinoid receptor 1 (CB1) incorporating unnatural amino-acid TCO*A before and after SPIEDAC reaction with methyl tetrazine dyes Me-Tet-ATTO488 or Me-Tet-ATTO655. **B.** Confocal micrographs showing CB1-F180 or CB2-S29 and R-GECO co-transfected HeLa Kyoto cells after receptor’s labelling with Me-Tet-ATTO488 (1 μM) for 30 min. **C.** Comparison of maximum fluorescence increase of R-GECO in CB1-F180-transfected HeLa Kyoto after treatment with WIN55,212-2 (10 μM, left graph) and with ATP (100 μM, right graph) under different experimental conditions. **D.** Comparison of maximum fluorescence increase of R-GECO in CB2-S29-transfected HeLa Kyoto after treatment with WIN55,212-2 or HU308 (both at 10 μM, left graphs) and with ATP (100 μM, right graphs) under different experimental conditions.

For studying CB2 signaling, HeLa cells were transfected with the CB2-S29 mutant. We measured [Ca^2+^]_i_ transients after receptor activation by the unspecific agonist WIN55,212-2 (F/F_0_=2.294±0.099, n=80, Figure 2D). Pre-treatment with the CB2-specific reverse agonist AM630 blocked the agonist-mediated calcium response in a concentration-dependent manner (F/F_0_=1.947±0.049, n=72, and F/F_0_=1.473±0.055, n=58 at 10 μM and 40 μM AM630, respectively. The CB2-specific agonist HU308 also triggered a calcium transient (F/F_0_=1.432±0.021, n=140) that was sensitive to the presence of AM630 (20 μM, F/F_0_=1.149±0.003, n=75). Unlike in CB1-expressing cells, we observed that both HU308- and WIN55,212-2-mediated calcium transients were blocked by pre-treating the cells with the PKA inhibitor H89 (F/F_0_=1.316±0.027, n=108 and F/F_0_=1.134±0.005, n=108 for HU308 and WIN55,212-2, respectively). This suggested CB2-specific involvement of PKA in calcium release. While H89 and AM630 separately blocked the CB agonist-mediated calcium transient, the ATP-mediated calcium transient was enhanced when the cells were treated with the CB2 reverse agonist but not with H89 (F/F_0_=3.397±0.130, n=72; F/F_0_=4.256±0.128, n=58 and F/F_0_=2.950±0.102, n=108, cells treated with 10 μM or 40 μM AM630 and 10 μM H89, respectively, Figure 2D). These results indicate that both reverse agonist-bound CB1 and CB2 receptors potentiate the cell response to the presence of extracellular ATP.

We previously showed that co-expressed CB1 and CB2 receptors were acting as a pair in regulating intracellular calcium oscillation in mouse β-cells as well as in human pancreatic islets of Langerhans.^35^ Endogenously co-expressed cannabinoid receptors 1 and 2 were suggested to act together as a switch participating in the regulation of cell excitability and insulin secretion. While CB1 activation could initiate calcium oscillations in silent cells, CB2 activation would damped oscillations and impact insulin secretion. In ß-cells, PKA, Ca^2+^ and cAMP exist in an oscillatory circuit characterized by a high degree of feedback.^50^. PKA activity has been demonstrated to be essential for this oscillatory circuit and is capable of not only initiating the signaling oscillations but also modulating their frequency.^51^

In our reconstituted model, the observation that PKA affected CB1 and CB2 signaling differently led us to interrogate the receptor function in regulating the activity of adenylate cyclase (AC) and hence endogenous cAMP levels. For this, we co-transfected CB1-F180 or CB2-S29 with a genetically encoded EPAC-based cAMP-sensor and measured changes in cAMP levels after stimulating the two GPCRs. A positive post-translational labeling of cannabinoid receptor 1 or 2 with cell-permeant far-red fluorescent ATTO 655 MeTet (Figure 2A) allowed us to distinguish between transfected and non-transfected cells in the same well (respectively noted * and #, Figure 3A). Single cell analysis revealed significantly different effects on cAMP levels between transfected (*i.e.*, labelled) and non-transfected (*i.e.*, unlabelled) cells present within the same field of view. Importantly, we did not observe a significant difference on cAMP levels between non-labelled cells (#) and wild type cells (1.254±0.007, n=178 and 1.296±0.009, n=76, respectively, Figures 3B, 3C and 3D). By treating the CB1-F180- or CB2-S29-expressing cells with WIN55,212-2 or with HU308, respectively (both at 10 μM) followed by a stimulation with forskolin (50 μM), we observed largely different effects on [cAMP]_i_ levels. While HU308 in CB2-S29-transfected cells did not directly affect the sensor’s FRET ratio, treating CB1-F180 cells with WIN55,212-2 induced a moderate increase in [cAMP]_i_ levels. WIN55,212-2 was unable of inducing a comparable increase in [cAMP]_i_ in CB2-S29-expressing cells, demonstrating the specificity of CB1 in mediating this effect (Figure 3E, F and S3E). Moreover, these observations reflected recent findings showing that CB1-mediated increase in [cAMP]_i_ levels correlated with reduced Gi/o function.^52^

**Figure 3.**
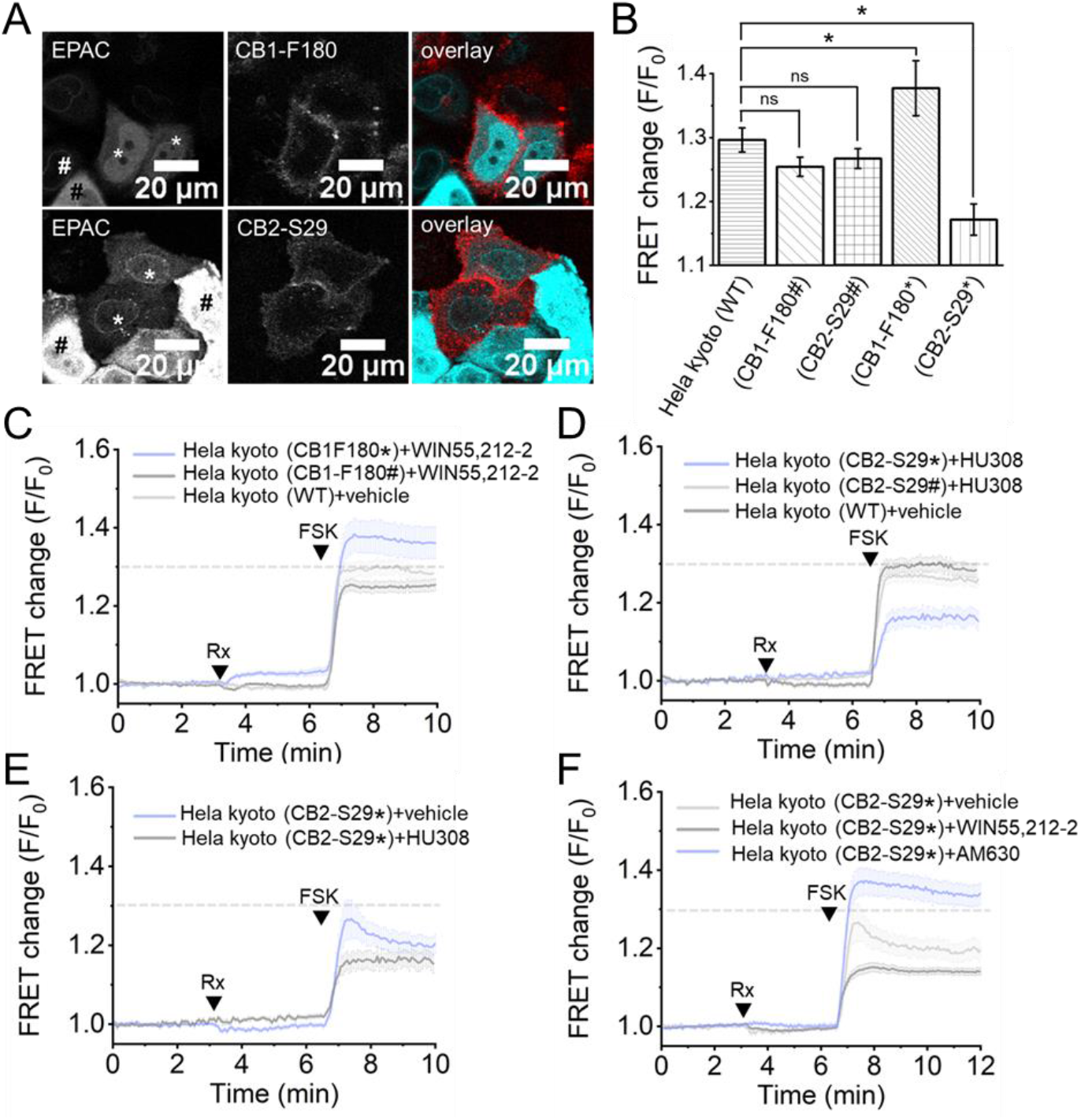
**A.** Confocal micrographs showing HeLa Kyoto cells co-transfected with CB1-F180 or CB2-S29 and the EPAC-based FRET sensor. Receptors were labelled with Me-Tet-ATTO655 (1 μM) for 30 min. Note that non-transfected cells in the field of view (#) served as controls. **B.** Bar graphs showing FRET changes of the EPAC-based FRET sensor after forskolin stimulation (50 μM) in wild type (WT) versus CB1-F180 or CB2-S29 expressing cells tagged with Me-Tet-ATTO655 (*) or non-labelled cells (#). WT/CB2-S29 p-value < 0.001, WT/CB1-F180 p-value < 0.05 **C.** Average of 45 and 178 traces showing FRET changes of the EPAC-based FRET sensor in CB1-F180-transfected HeLa (*) and (#), respectively, after treatment with WIN55,212-2 (10 μM) followed by forskolin (FSK, 50 μM). **D.** Average of 16 and 91 traces showing FRET changes of EPAC-based FRET sensor in CB2-S29-transfected HeLa cells (*) and (#), respectively, after treatment with HU308 (10 μM) followed by forskolin (FSK, 50 μM). **E.** Average of 16 and 21 traces showing differences in FRET changes in CB2-S29-transfected HeLa Kyoto (*) after treatment with HU308 (10 μM) or vehicle, followed by forskolin stimulation (FSK, 50 μM). **F.** Average of 16, 27 and 21 traces showing differences in FRET changes in CB2-S29-transfected HeLa Kyoto after treatment with HU308, AM630 (both at 10 μM), or vehicle, respectively, followed by forskolin stimulation (FSK, 50 μM).

The difference in CB1-F180- and CB2-S29-mediated signaling was much more pronounced after forskolin stimulation. Here, CB1-F180-transfected cells showed moderately higher [cAMP]_i_ levels (1.377±0.022, n=45, Figure 3B and C) than non-transfected (#) or wild types cells. Conversely, forskolin-stimulated CB2-S29-transfected cells (*) showed significantly lower [cAMP]_i_ levels than the non-transfected (#) or wild types cells (1.15714±0.024, n=16, Figure 3B and D). Control experiments performed with native and untreated CB1 or CB2 receptors confirmed our preliminary observations (Figure S3A). These results suggest that CB1 and CB2 can affect the AC activity in absence of pharmacological activation with synthetic agonists. Moreover, in CB1-F180 and CB2-S29-expressing cells, the FRET change of the EPAC-based sensor was comparable to control experiments where the cells were transfected with the native CB1 and CB2 receptors under the same conditions (Figure S3A, C and D).

Both cannabinoid receptors 1 and 2 are known to be Gi/o coupled,^53,54^ leading to reduced cAMP levels by inhibiting adenylyl cyclase (AC) activity and subsequently cAMP-dependent protein kinase activity.^18^ Therefore, we were interested to compare agonist-mediated activation of CB1 or CB2 to non-stimulated cells. We hypothesize that CB1 and CB2 activation leads to an inhibition in AC activity. Indeed, WIN55,212-2 stimulation of CB1-transfected cells marginally decreased forskolin-stimulated [cAMP]_i_ levels (Figure S3B) and treatment with the inverse agonist rimonabant significantly increased forskolin-stimulated [cAMP]_i_ levels (Figure S3B). Similarly, the treatment of CB2-S29 transfected cells with synthetic agonists (*i.e.*, HU308 or WIN55,212-2, Figure 3E and F) also lowered forskolin-stimulated [cAMP]_i_ levels compared to cells treated with vehicle only, while CB2-S29 transfected cells treated with the inverse agonist AM630 showed significantly higher levels of [cAMP]_i_ after stimulation compared to control (Figure 3F and S3F).

The different functions of CB1 and CB2 in regulating stimulated [cAMP]_i_ levels led us to investigate the effect of their co-expression on [Ca^2+^]_i_ levels. As discussed above, the activation of individually transfected CB1 or CB2 leads to a transient elevation of [Ca^2+^]_i_ levels. Moreover, the CB1- and CB2-mediated elevation of [Ca^2+^]_i_ was prevented by pre-treating the cells with the respective reverse agonists. In β-cells however, while both CB1 and CB2 receptors are endogenously present, the transient elevation of [Ca^2+^]_i_ levels mediated by WIN55,212-2 was completely blocked after pre-treatment with rimonabant. This observation suggested the specificity of CB1 in mediating this effect. Therefore, we investigated if co-expressing both CB1 and CB2 in our reconstituted model would give similar results.

For this, we co-expressed CB1-GFP and CB2-S29 as well as the calcium sensor R-GECO in HeLa cells. The tagging of CB2-S29 with Me-Tet ATTO 655 demonstrated the co-localization of both receptors at the plasma membrane as well as on intracellular vesicles (Figure 4A). Under these conditions, the addition of WIN55,212-2 triggered a transient increase of [Ca^2+^]_i_ levels (F/F_0_=2.187±0.061, n=223). Similar to the β-cell system, we observed that the formation of the transient was completely blocked by pre-incubating the cells with rimonabant (F/F_0_=1.299±0.034, n=129, Figure 4B and C). Conversely, HU308 alone did not elicit a calcium transient (F/F_0_=1.119±0.004, n=193). Finally, the treatment with WIN55,212-2 and ATP showed weaker effects on [Ca^2+^]_i_ levels in the co-transfected model than in the CB1-expressing cells. Taken together, these observations suggest that 1) CB2 expression can affect CB1 receptor signaling at the calcium level and 2) co-expressing both CB1 and CB2 decreases the cell sensibility to extracellular ATP.

**Figure 4.**
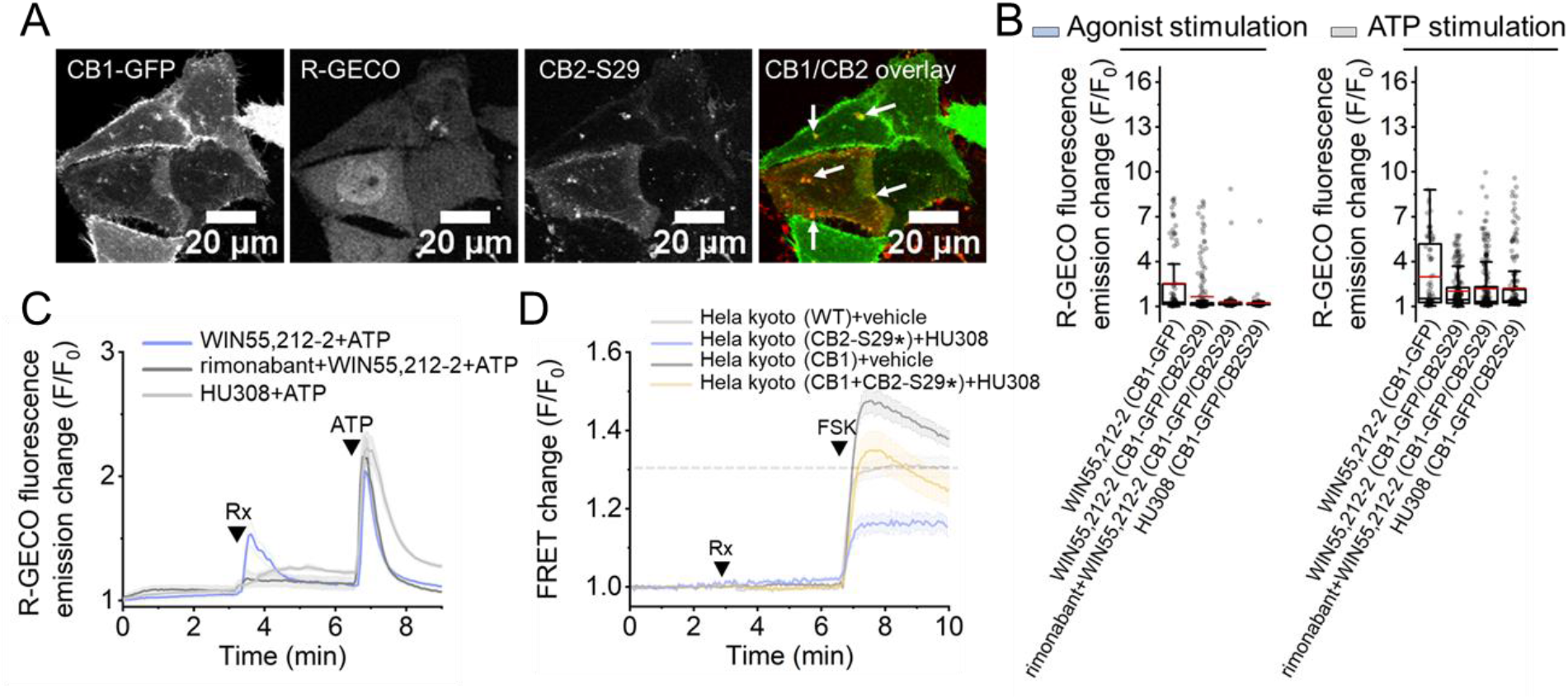
**A.** Confocal micrograph showing the co-transfection of CB1-GFP, R-GECO and CB2-S29 in HeLa Kyoto cells. CB2-S29 was labelled with Me-Tet-ATTO655 (1 μM). Arrows show co-localization of CB1-GFP and tagged CB2-S29. **B.** Graph showing the maximum fluorescence increase F/F_0_ of R-GECO after treatment with agonist in presence or absence of inverse agonist (10 μM each, left panel) and after treatment with ATP (100 μM, right panel). **C.** Corresponding averaged calcium traces. **D.** FRET changes of EPAC-based FRET sensor in CB1/CB2-S29 co-transfected HeLa Kyoto after treatment with vehicle or HU308 (10 μM) and forskolin (FSK, 50 μM).

Finally, we investigated whether these effects on calcium would correlate with the AC activity. For this, the changes in [cAMP]_i_ levels in response to the treatment with HU308 (10 μM) followed by forskolin were compared between CB2-S29- and CB1/CB2-S29 co-expressing cells. Remarkably, the biphasic response of CB1/CB2-S29 co-transfected cells after forskolin stimulation showed additive –yet opposing– effects of the two receptors (Figure 4D). This result indicates that CB1 and CB2 are acting via the same effectors but independent of each other.

## Discussion and conclusions

While CB1 and CB2 share only 44% structural homology, both receptors are fully activated by 2-AG and associate with the same G-proteins. Surprisingly, antagonist signaling outcomes resulting from the selective activation of CB1 or CB2 have suggested opposing roles of the two receptors in different cell types and tissues. The structural comparison between the antagonist-bound CB2 and the agonist-bound CB1 that shows superimposable structures of the two receptors under these conformations supports these observations.^12^ In this study, we developed a simple cell system model to interrogate the roles of CB1 and CB2 co-expression in regulating calcium and cAMP levels. We applied amber codon suppression in live cells to incorporate a single *trans*-cyclooctene (TCO) bearing amino acid in one of the extracellular loops of CB1 or CB2, followed by fluorescent labeling *via* copper-free click chemistry. We simultaneously used genetically encoded biosensors to investigate the roles of CB1 and/or CB2 in regulating intracellular calcium ([Ca^2+^]_i_) and cAMP ([cAMP]_i_) levels. By incorporating a single unnatural amino-acid for receptor labeling, a palette of tetrazine-bearing fluorophores can be used to label the CB-receptors without conflicting with the fluorescent reporters of the biosensors employed during an experiment. Furthermore, we demonstrated the feasibility of incorporating a single unnatural amino-acid in CB1 or CB2 at a specific position without impeding the signaling ability of the receptors on calcium and cAMP –as the reconstituted model mimics the cell behavior of endogenous systems. This is due to the small size of the fluorescent tag that induces minimal changes in the receptors structure.

By using this strategy, we showed that both the agonist-mediated activation of tagged-CB1 or - CB2 can transiently elevate [Ca^2+^]_i_ levels. However, we demonstrated that when the two receptors co-expressed in the same cell, CB2 no longer signaled through calcium although CB1-mediated transient elevation of [Ca^2+^]_i_ levels remained unaffected. We also measured the effects of CB1 and/or CB2 in regulating stimulated-adenylate cyclase activity in presence or absence of synthetic agonists. We found that the expression of CB1 increased forskolin-induced [cAMP]_i_ levels compared to non-transfected cells. Conversely, CB2 expression decreased stimulated [cAMP]_i_ levels under the same conditions. Finally, co-expressed CB1 and CB2 receptors showed additive yet opposing effects on stimulated [cAMP]_i_ levels. Our key findings indicate that the expression of CB1 and CB2 have an opposing effect in regulating [cAMP]_i_ level upon cell stimulation with forskolin, while the stimulation of these receptors by agonists have similar effects. As a result, the effects of CB1 and CB2 on modulating cAMP levels in the HeLa model system is not only mediated by stimulation with synthetic agonists, but also by the sole expression of the receptors (Figure 3 and S3). Further investigations are now needed to evaluate the role of endogenous endocannabinoids in our cell model, as well as their contribution to the constitutive activation level of the receptors.

Further experiments will also be required to investigate if this effect is mediated by direct interactions between CB1 and CB2 (*i.e.*, namely by forming heteromers),^55^ or via interaction with other proteins. For example, experimental data have shown that most class A rhodopsin-like G-protein coupled receptors could form a mixture of monomers and homodimers/oligomers.^56^ Several reports show that CB1 is engaging in the formation of protein complexes with other GPCRs^57^ (*i.e.* adenosine A2A receptor,^58^ dopamine D2 receptor,^59^ opioid μ and δ receptors,^60,61^ orexin OX1 receptor,^62^ angiotensin AT1 receptor,^63^ adrenergic β2 receptor,^64^ 5HT2A serotonin receptor^65^ and CB2 receptor^55^). In the other hand, CB2 has been reported to associate with other GPCRs such as GPR55 and CXCR4,^66,67^ as well as the receptor tyrosine kinase (RTK) HER2.^68^ These interactions may also shape the receptors response to pharmacological activation. Interestingly, most of these protein complexes are reported to form at the cell surface. Similarly, most studies on CB1 and CB2 have focused on their localization to the plasma membrane. However, evidence suggests that fully functional CB1 and CB2 receptors are also located in endomembranes.^69,70^ For example, intracellular CB1 was suggested to exhibit different functions distinct from plasma membrane CB1 receptors.^71^ While CB1 mediates the activation of G-protein gated inwardly rectifying potassium (GIRK) channels at the cell surface, the activation of mitochondrial CB1 (*i.e.*, mtCB1) decreased complex I enzymatic activity and respiration in neuronal mitochondria.^72^

For future studies, it will be interesting to probe the different roles of CB1 or CB2 activation depending on where the receptors accumulate inside the cell. We can already hypothesize that while intracellular receptors activation will affect endolysosomal calcium levels as well as mitochondrial activity, plasma membrane-bound receptors will predominantly act on cAMP levels via Gi/o signaling. We previously described a method for increasing intracellular endocannabinoid levels with cg2-AG.^34^ Additional chemical modifications of cg2-AG such as adding sulfonated groups to the coumarin cage would be a promising strategy allowing for the exclusive photo-release of the endocannabinoid at the plasma membrane.^73^ Other chemical modifications that will drive the accumulation of cg2-AG to specific organelles such as mitochondria or lysosomes before uncaging can be found in this review.^74^

Interestingly, 2-AG has a three-fold stronger binding affinity towards CB1 than CB2.^75^ Because 2-AG is a degradation product of diacylglycerols, we can assume than no cell can avoid making 2-AG. Therefore, in a localized subcellular environment where both CB1 and CB2 co-express, an increase of 2-AG levels is likely to activate the two receptors independently and sequentially. As 2-AG activates both receptors, it can be anticipated that 2-AG levels also coordinate interactions between CB1, CB2 and other proteins. Hence, there is a need in the endocannabinoid research community for studying cannabinoid receptors interactions with other proteins, beyond the established role of heterotrimeric G-proteins. Some recent studies suggested the biological relevance of cannabinoid receptors forming heteromers in human diseases such as diabetes,^76^ obesity,^77^ cancer^78^ as well as neurodegenerative disorders.^79^ The prevalence, the biological functions and the composition of cannabinoid receptor heteromers in living cells remain obscure and needs further investigations. While our results further demonstrate that cannabinoid receptors need to be jointly investigated, we hypothesize that the synergistic activity between cannabinoid receptors may play crucial roles in cells and tissues that can be therapeutically leveraged for the treatment of human diseases.

---

**Figure S1.**
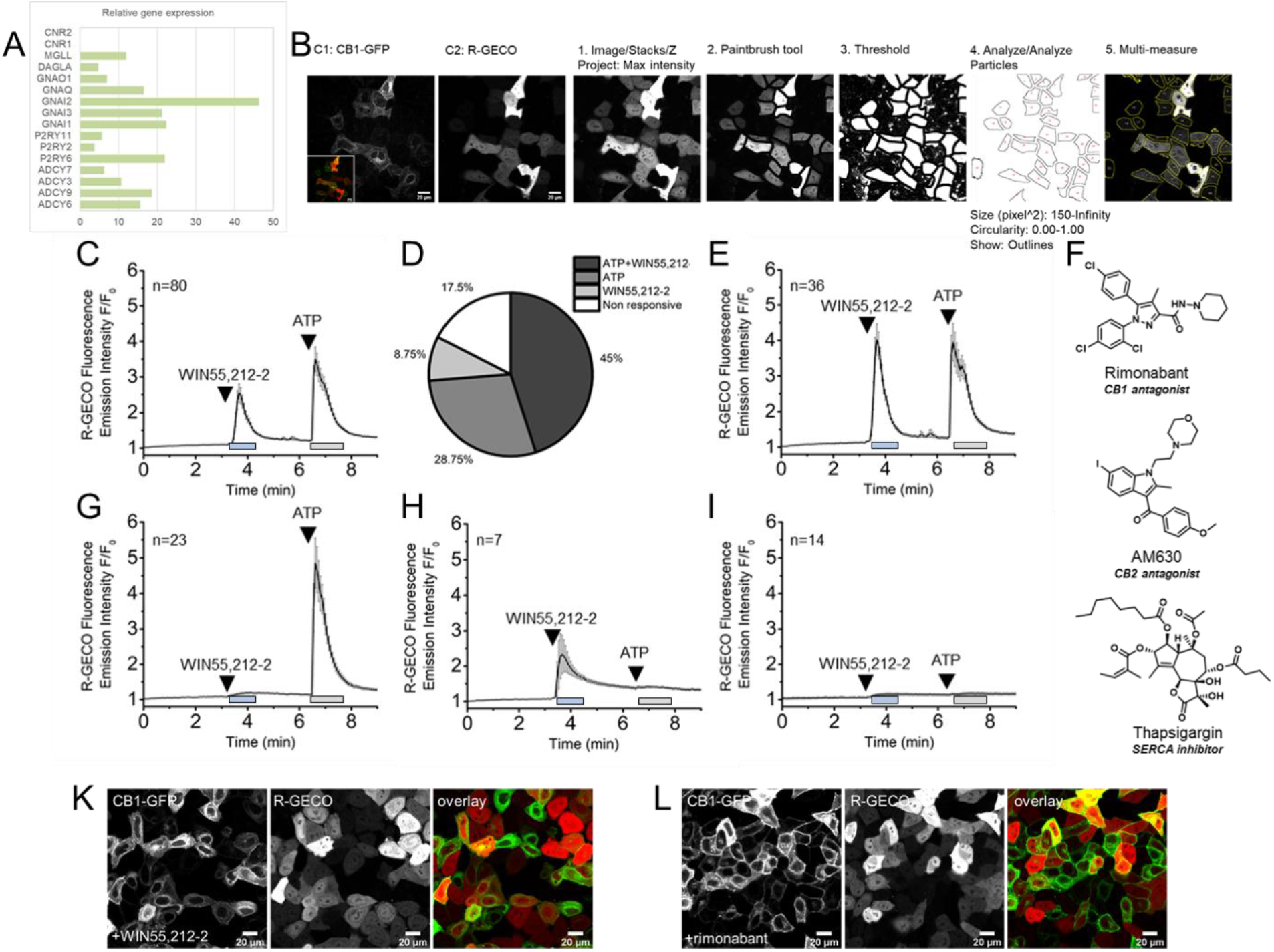
**A.** Relative genes expression in HeLa Kyoto cells of adenylate cyclase (ADCY) and purinergic receptor (P2RY) isoforms, as well as Gi (GNAI), Go (GNA0), Gq (GNAQ), MAGL (MGLL), DAGLα (DAGLA), CB1 (CNR1) and CB2 (CNR2) receptors. **B.** Step-by-step cell segmentation procedure for single-cell data analysis of calcium imaging experiments. **C.** Averaged R-GECO fluorescence emission intensity changes (F/F_0_) of CB1-GFP-transfected HeLa Kyoto (n=80) upon treatment with WIN55,212-2 (10 μM) and with ATP (50 μM). **D.** Pie-chart showing the repartition of the different cell populations responding to treatments with CB1 agonist and ATP. **E.** Cell population responding to both WIN55,212-2 and ATP. **F.** Chemical structures of the CB1 inverse agonist rimonabant, CB2 inverse agonist AM630 and SERCA inhibitor thapsigargin. **G.** Cell population responding to ATP only. **H.** Cell population responding to WIN55,212-2 only. **I.** Non responsive cells. Confocal micrographs (20x) showing CB1-GFP and R-GECO co-transfected cells after 3 hr treatment with 10 μM of WIN55,212-2 (**K**) or with 10 μM of rimonabant (**L**).

**Figure S3.**
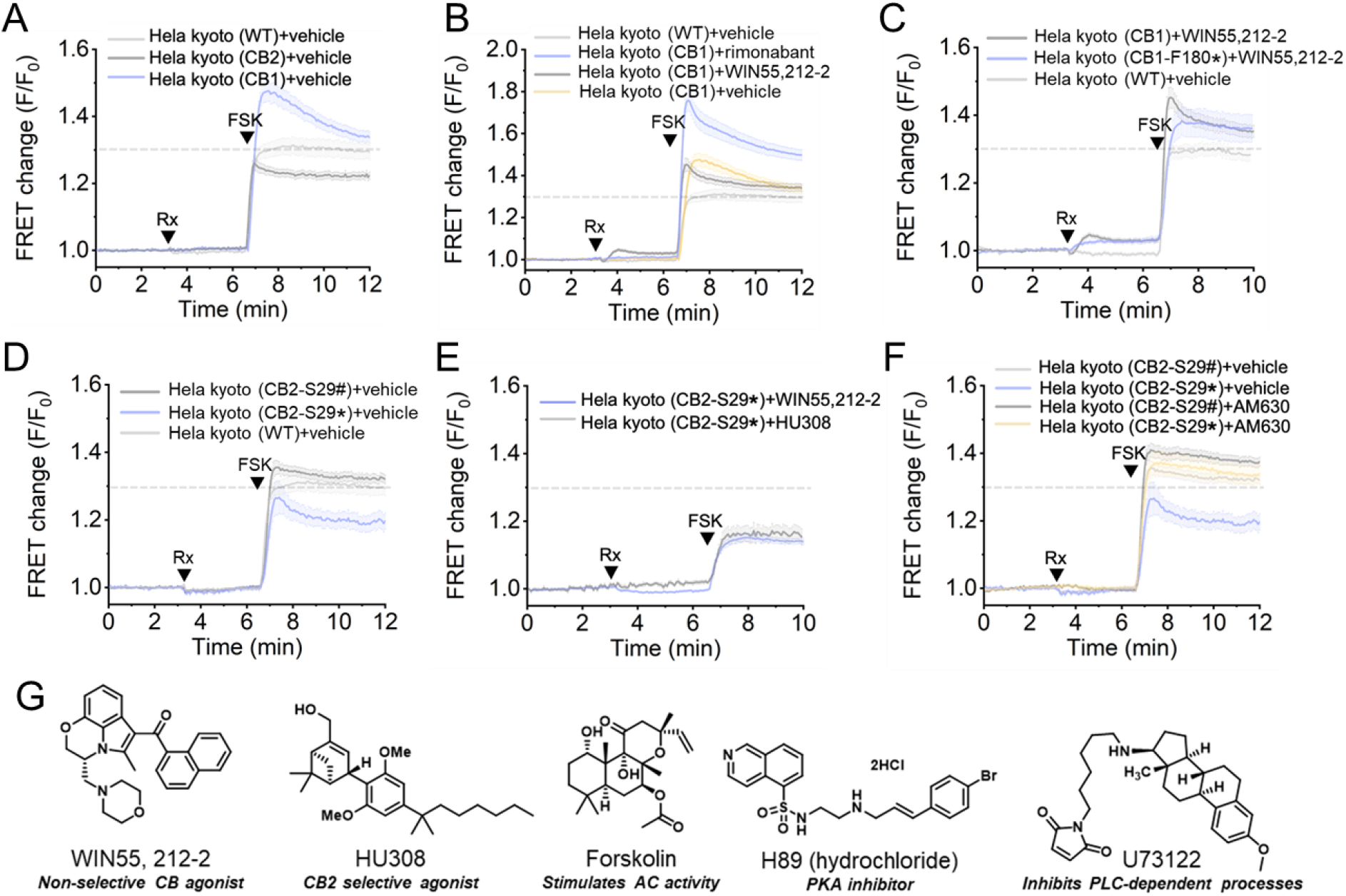
**A.** Averaged traces showing differences in FRET changes of the EPAC-based FRET sensor between wild types (n=76), CB1 (n=56) or CB2 (n=65) transfected HeLa Kyoto cells after treatment with vehicle followed by forskolin (FSK, 50 μM). **B.** Averaged traces showing differences in FRET changes between wild types (n=70) and CB1-transfected HeLa Kyoto after treatment with WIN55,212-2, rimonabant (both at 10 μM, n=70 and n=49, respectively) or vehicle (n=76), followed by forskolin stimulation (FSK, 50 μM). **C.** Comparison between non-mutated CB1 (n=70) and CB1-F180* (n=45) upon treatment with WIN55,212-2 (10 μM) and with forskolin (FSK, 50 μM). **D.** FRET changes of CB2-S29* (n=21) and CB2-S29# (n=59) compared to wild type cells (n=76) after forskolin stimulation (FSK, 50 μM). **E.** FRET changes of CB2-S29* cells treated with WIN55,212-2 (10 μM, n=21) or with HU308 (10 μM, n=16) and with forskolin (FSK, 50 μM). **F.** FRET changes of CB2-S29* (n=27) and CB2-S29# (n=115) treated with AM630 (10 μM) or vehicle (n=21 and n=59, respectively) and with forskolin (FSK, 50 μM). **G.** Chemical structures of WIN55,212-2, HU308, forskolin, H89 and U73122.

**Table S1.** Residues replaced by TCO*A lysine amino-acid in CB1 or CB2 that demonstrated a successful labelling with tetrazine dyes (+), or not (−).

|  | Labeling |  |
| --- | --- | --- |
|  | + | - |
| **CB1** | F109 | M110 |
|  | F180 | L112 |
|  | K184 | H182 |
|  | K371 | K260 |
|  |  | E262 |
|  |  | D267 |
|  |  | L271 |
| **CB2** | I27 | T173 |
|  | S29 | E181 |
|  | H98 | L185 |
|  | V100 | T272 |
|  |  | S274 |

## STAR METHODS

### RESOURCE AVAILABILITY

#### Lead Contact

Further information and requests for resources and reagents should be directed to and will be fulfilled by the Lead Contact, Aurélien Laguerre.

#### Materials Availability

This study did not generate new unique reagents.

#### Data and Code Availability

This study did not generate any unique datasets or code.

### EXPERIMENTAL MODEL AND SUBJECT DETAILS

The HeLa Kyoto cell line (RRID:CVCL_1922, female) was kindly provided by R. Pepperkok (European Molecular Biology Laboratory, Germany). HeLa Kyoto (passage 15-35) were grown in 4.5g/L glucose DMEM (Life Technologies, 41965-039) supplied with 10 % fetal bovine serum (Life Technologies, 10270098).

### METHOD DETAILS

#### General

All chemicals were obtained from commercial sources (Acros, Sigma-Aldrich, Tocris, TCI, Cayman, Alfa Aesar, Atto-tec or Merck) and were used without further purification unless otherwise specified. Rimonabant (5-(4-chlorophenyl)-1-(2,4-dichlorophenyl)-4-methyl-N-(piperidin-1-yl)-1H-pyrazole-3-carboxamide), AM630 ((6-iodo-2-methyl-1-(2-morpholinoethyl)-1H-indol-3-yl)(4-methoxyphenyl) methanone, HU308 (4-[4-(1,1-dimethylheptyl)-2,6-dimethoxyphenyl]-6,6-dimethyl-bicyclo[3.1.1]hept-2-ene-2-methanol, H-89 hydrochloride (N-[2-[[3-(4-bromophenyl)-2-propen-1-yl]amino]ethyl]5-isoquinolinesulfonamide, dihydrochloride, Forskolin (5-(acetyloxy)-3-ethenyldodecahydro-6,10,10b-trihydroxy-3,4a,7,7,10a-pentamethyl-(3R,4aR,5S,6S,6aS,10S,10aR,10bS)-1H-naphtho[2,1-b]pyran-1-one) and WIN55, 212-2 ([(11R)-2-methyl-11-(morpholin-4-ylmethyl)-9-oxa1-azatricyclo[6.3.1.04,12]dodeca-2,4(12),5,7-tetraen-3-yl]-naphthalen-1-ylmethanone) from Cayman Chemical were dissolved in dimethylsulfoxide (DMSO) to a stock concentration of 10 mM. Thapsigargin ((3S,3aS,4R,6R,7S,8R)-6-acetoxy-4-(butyryloxy)-3,3a-dihydroxy-3,6,9-trimethyl-8-(((Z)-2-methylbut-2-enoyl)oxy)-2-oxo-2,3,3a,4,5,6,6a,7,8,9b-decahydro-1H-cyclopenta[e]azulen-7-yl octanoate) from sigma was dissolved in DMSO to a stock concentration of 5 mM. U73122 (1-[6-[[(17β)-3-methoxyestra-1,3,5(10)-trien-17-yl]amino]hexyl]-1H-pyrrole-2,5-dione) from Cayman was dissolved in DMSO to a stock concentration of 10 mM. ATP (adenosine 5’-triphosphate disodium salt hydrate) from TCI was freshly dissolved in DMSO to a concentration of 10 mM. Atto488 Me-Tetrazine from Atto-Tec was dissolved in DMSO to a stock concentration of 1 mM. All chemicals were administrated to cells with a DMSO concentration lower or equal to 0.1 %.

#### Genetic code expansion and SPIEDAC tagging

Cells were seeded in eight-well Lab-Tek microscope dishes for 24 hr (to reach 60-70 % confluence) before transfection. After 24 hr, 200 ng of hMbPylRS-4xU6M15 (Addgene, #105830) and 200 ng of the respective amber construct were premixed in 20 μL of DMEM. 1.5 μL of Lipofectamine 2000 (Life Technologies, 11668030) in 20 μL of DMEM was then added to the DNA premix and incubated for 20 min at RT before being added to the wells. Shortly after the transfection mixture was added to cells, 100 μM of the ncAA TCOA*K was added from a 100 mM stock solution in 0.1 M NaOH. After overnight incubation the transfection medium was replaced with fresh full growth medium. 30 min before imaging, cells were washed two times with DMEM (without FBS) and incubated for 20 min with 1 μM of Me-Tet Atto488 from a 1 mM stock solution in DMSO. After 20 min cells were washed with imaging medium (Invitrogen, A14291DJ) four times before imaging.

#### Calcium imaging experiments

Cells were seeded in eight-well Lab-Tek microscope dishes for 24 hr (to reach 60-70 % confluence) before transfection. For imaging of CB1-GFP transfected cells, 100 ng of CB1-GFP and 100 ng of R-GECO (Addgene #32444) were mixed with 1 μL of lipofectamine 2000 transfection reagent. For imaging of cells transfected with CB1-F180 or CB2-S29, the experimental protocol described in the genetic code expansion section was followed with an addition of 200 ng of R-GECO. For imaging of cells co-transfected with CB1-GFP and CB2-S29, the experimental protocol described in the genetic code expansion section was followed with an addition of 100 ng of R-GECO and 100 ng of CB1-GFP. For all of the above mixes, DNAs and lipofectamine were separately premixed in 20 μL of DMEM then mixed together and incubated for 20 min before being added to each well of the eight well Lab-Tek containing 200 μL of DMEM 4.5g/L glucose supplemented with 10% FBS. Cells were imaged at 37°C in imaging buffer. Imaging was performed on a dual scanner confocal microscope Olympus Fluoview 1200, with a 63x (oil) objectives. The R-GECO sensor was imaged using a 559 nm laser (LP, 1.0 %) and a 643/50 emission filter. Fluctuations of [Ca^2+^]_i_ were monitored through excitation at 559 nm and emission above 600 nm (F/F_0_) on the confocal microscope.

#### Trafficking experiments

Cells were seeded in eight-well Lab-Tek microscope dishes for 24 hr before transfection. 100 ng of CB1-GFP and 100 ng of Rab5-BFP (Addgene #49147) were mixed with 1 μL of lipofectamine 2000 following transfection method previously described then added to the wells. 48 hr after the transfection, cells were incubated with 10 μM of WIN55,212-22 or 10 μM of rimonabant for 3 hr. Cells were imaged at 37°C in imaging buffer. Imaging was performed on a dual scanner confocal microscope Olympus Fluoview 1200, with a 63x (oil) objectives.

#### EPAC-based sensor imaging experiments

Cells were seeded in eight-well Lab-Tek microscope dishes for 24 hr (to reach 60-70 % confluence) before transfection. 200 ng of the EPAC Sensor (Addgene #61622) were mixed with 1 μL of lipofectamine 2000 following the transfection method previously described. After overnight incubation the transfection medium was replaced with fresh full growth medium. 24 hr after the first transfection, native CB1 (200 ng) or CB2 (200 ng) were mixed with 1 μL of lipofectamine 2000 and added to cells. For CB1-F180 or CB2-S29, the experimental protocol described in the genetic code expansion section was used after the cells were first transfected with the EPAC sensor. Cells were imaged at 37°C in imaging buffer. Imaging was performed on a dual scanner confocal microscope Olympus Fluoview 1200, with a 63x (oil) objectives. The EPAC FRET sensor was imaged using a 440 nm laser (LP, 1,0%) and the signal collected in the CFP/YFP emission channels.

#### Images analysis

All Images were analyzed on the FIJI software using the pipeline summarized in Figure S1B. Primarily, multi-channel images were separated into single channels and converted to 8-bit for calcium imaging or 32-bit for EPAC experiments. The time course experiment was duplicated and stacked using the Z project function (RFP channel for calcium imaging and CFP channel for EPAC imaging). Using the paintbrush tool set at 0, cells were manually delimited to achieve robust single cell segmentation. A mask of regions of interest was generated using the combination of the threshold and analyze particles tools (as depicted in Figure S1B). This ROI mask was then superimposed to the time course experiment and the multi-measure function was applied to it. From this stack, we extracted mean single cell values from the time course experiment. Those values were then exported to an excel files for further analysis.

#### Statistical analysis

All statistical comparisons were performed using Student t-test by Prism or Excel. Statistical details of each experiment can be found in the figures and figure legends. For all experiments, the number of cells and error bars (SEM) can be found in the results section and the respective figure legends. All imaging experiments were performed at least in biological triplicates, n indicating the total number of cells.

#### Primers

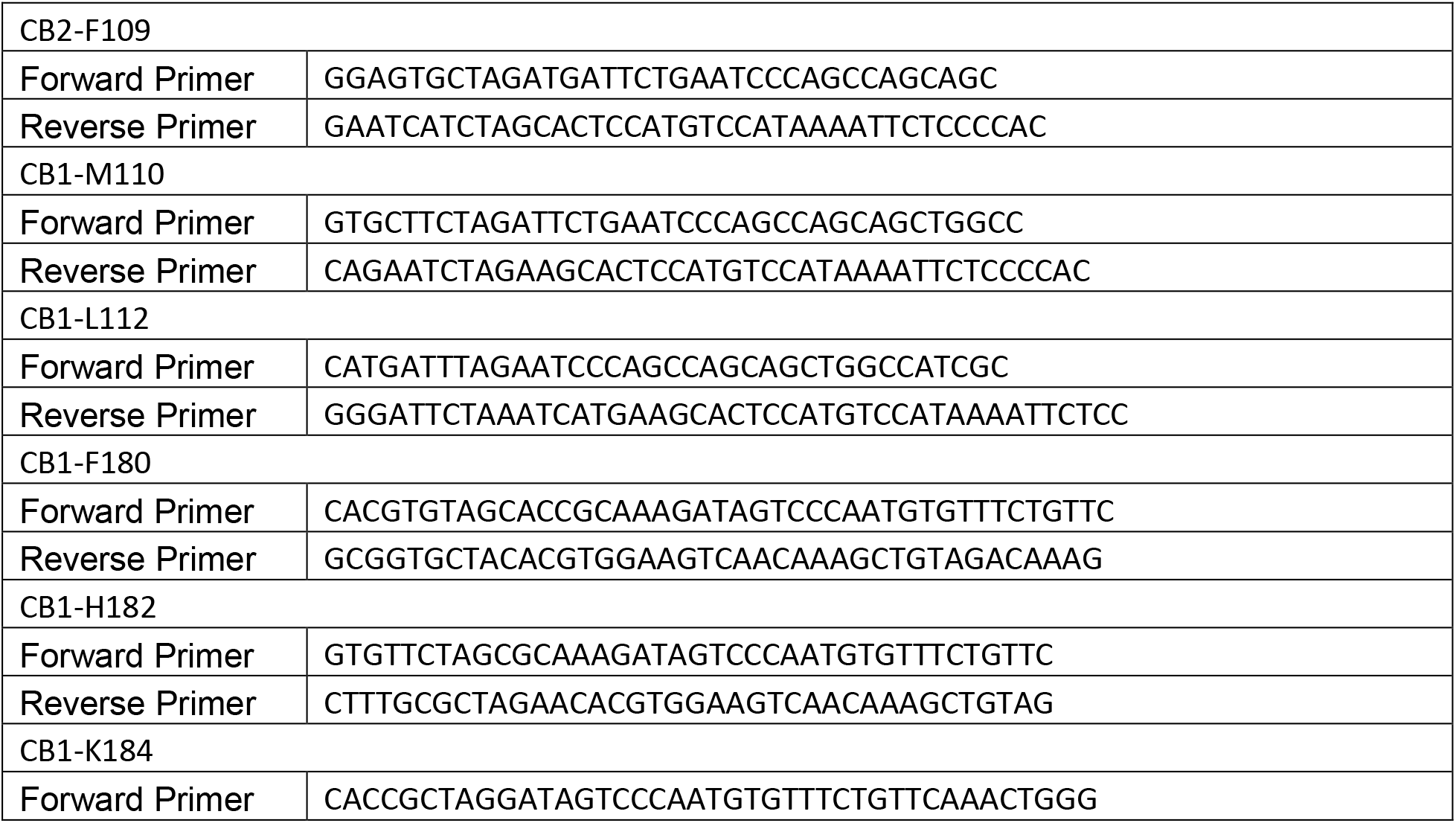
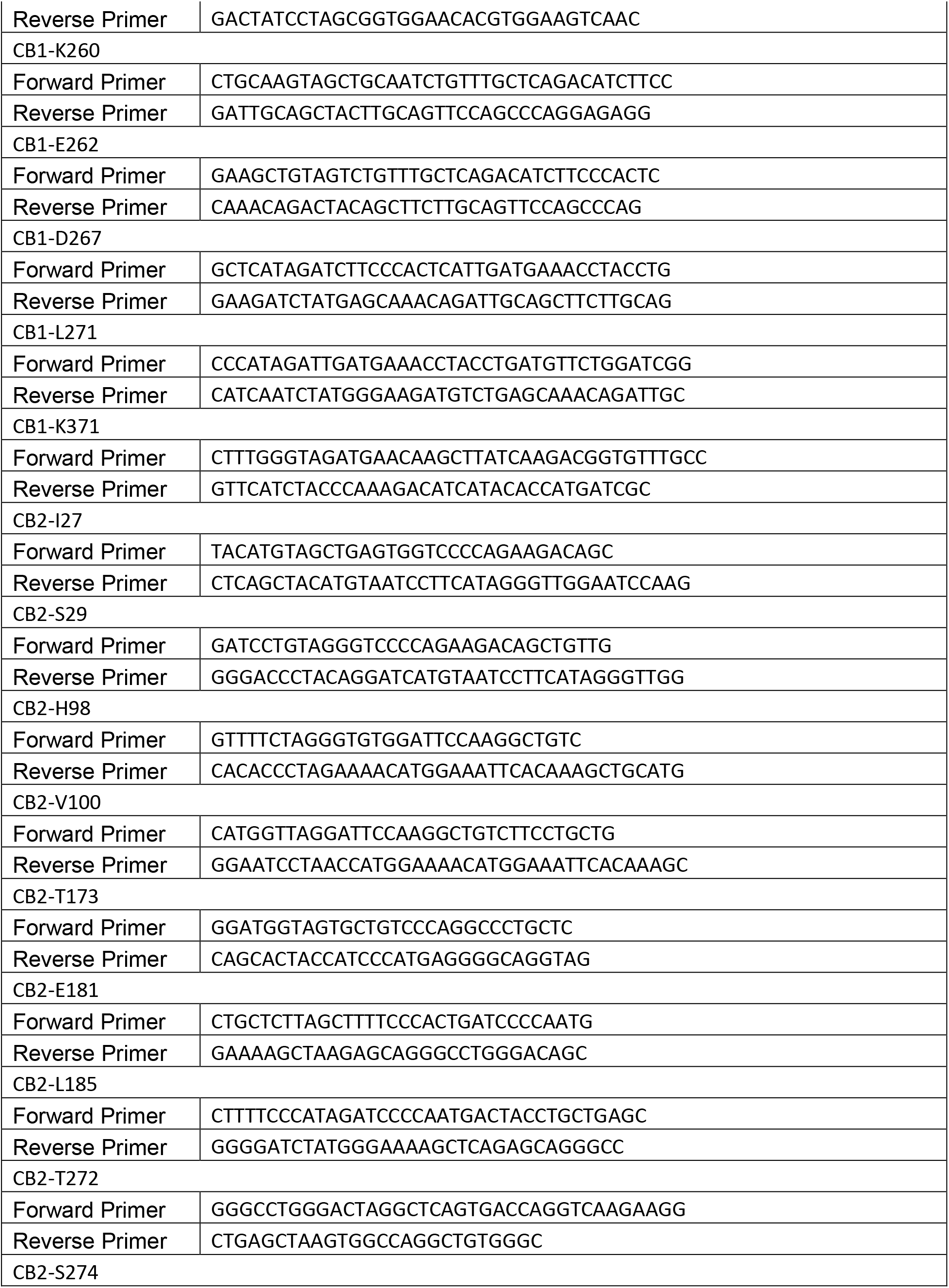
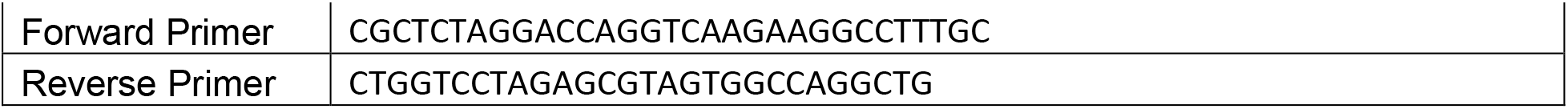

##### KEY RESOURCES TABLE

| REAGENT or RESOURCE | SOURCE | IDENTIFIER |
| --- | --- | --- |
| **Antibodies** |  |  |
| Rabbit polyclonal Cannabinoid Receptor I antibody | Abcam | Cat# ab3561 |
| Rabbit polyclonal Cannabinoid Receptor II antibody | Abcam | Cat# ab3561 |
| Goat anti-Rabbit IgG (H+L) Highly Cross-Adsorbed Secondary Antibody, Alexa Fluor Plus 488 | Invitrogen | Cat# A32731 |
| **Recombinant proteins** |  |  |
| hMbPyIRS-4xU6M15 | Addgene (Serfling et al,2017) | Cat# 105830 |
| CB1-GFP | Farrens Lab, OHSU | N/A |
| R-GECO | Addgene (Zhao et al, 2011) | Cat# 32444 |
| CB1 | Farrens Lab, OHSU | N/A |
| CB1-F180 | This study | N/A |
| CB2 | UMR cDNA RC (U Missouri) | Cat# AY242132 |
| CB2-S29 | This study | N/A |
| EPAC | Addgene (Dipilato et al, 2009) | Cat# 61622 |
| RAB5-BFP | Addgene (Friedman et al, 2013) | Cat# 49147 |
| **Chemicals, Peptides, and Recombinant Proteins** |  |  |
| Rimonabant | Cayman Chemical | Cat# 9000484 |
| AM630 | Cayman Chemical | Cat# 10006974 |
| WIN55,212-2 | Cayman Chemical | Cat# 10009023 |
| HU308 | Cayman Chemical | Cat# 90086 |
| H-89 hydrochloride | Cayman Chemical | Cat# 10010556 |
| U73122 | Cayman Chemical | Cat# 70740 |
| Forskolin | Cayman Chemical | Cat# 11018 |
| Thapsigargin | Sigma | Cat# T9033 |
| ATP | TCI | Cat# A0157 |
| Atto655 Me-Tetrazine | Atto-tec | Cat# SA04Z04 |
| Atto488 Me-Tetrazine | Atto-tec | Cat# 02102 |
| Trans-Cyclooct-2-en-L-Lysine | SiChem | Cat# SC-8008 |
| **Experimental Models: Cell Lines** |  |  |
| Kyoto Hela cells | (Hein M.Y et al, 2015) | RRID:CVCL_1922 |
| **Software and Algorithms** |  |  |
| FIJI | ImageJ | https://imagej.net/software/fiji/ |
| GraphPad Prism (RRID:SCR_002798) | Graphpad | https://www.graphpad.com/scientific-software/prism/ |

## ACKNOWLEDGEMENTS

The authors warmly thank Dr. Ana Kojic for many helpful discussions and critical reading of the manuscript. The authors also thank the Farrens and the Frank groups from OHSU for providing wild type CB1 and CB2 constructs. A.T., C.S. and A.L. acknowledge financial support by OHSU.

## AUTHOR CONTRIBUTIONS

A.T. performed cloning and imaging experiments, analyzed data and interpreted the results. C.S. interpreted the results and co-wrote the manuscript. A.L. designed the study, performed imaging experiments, analyzed data, interpreted the results and co-wrote the manuscript.

